# Theta and alpha power across fast and slow timescales in cognitive control

**DOI:** 10.1101/2020.08.21.259341

**Authors:** Pieter Huycke, Pieter Verbeke, C. Nico Boehler, Tom Verguts

## Abstract

Theta and alpha frequency neural oscillations are important for learning and cognitive control, but their exact role has remained obscure. In particular, it is unknown whether they operate at similar timescales, and whether they support different cognitive processes. We recorded EEG in 30 healthy human participants while they performed a learning task containing both novel (block-unique) and repeating stimuli. We investigated behavior and electrophysiology at both fast (i.e., within blocks) and slow (i.e., between blocks) time scales. Behaviorally, both response time and accuracy improved (resp. decrease and increase) over both fast and slow timescales. However, on the spectral level, theta power significantly decreased along the slow timescale, whereas alpha power instead significantly increased along the fast timescale. We thus demonstrate that theta and alpha both play a role during learning, but operate at different timescales. This result poses important empirical constraints for theories on learning, cognitive control, and neural oscillations.

## Introduction

In daily life, agents frequently have to remember, apply, and automatize novel stimulus-action mappings, a process which falls under the general umbrella of *cognitive control* (Botvinick et al., 2001; Ridderinkhof, 2004). An example is reacting to traffic signs. When you are new to driving a car, it might be challenging to bring the car to a halt when you see a nearby stop sign. But over time, with more practice, this process of seeing the sign and stopping becomes natural, and one can do it without giving it much thought (i.e., without cognitive control).

Previous research has started to identify the neural signatures of cognitive control. In the spectral domain, theta band (4 – 8 Hz) oscillations were originally observed in the hippocampus (Buzsáki, 2002; O’Keefe, 1993; Vertes & Kocsis, 1997) and considered to be crucial for memory formation. More recently, theta was also observed in structures of the cerebral cortex, and became intensively studied in the cognitive-control literature (Cavanagh & Frank, 2014). These theta oscillations are thought to be generated in the frontal-medial (FM) cortex of the human brain (Anguera et al., 2013; Enriquez-Geppert et al., 2014; Sauseng et al., 2019). In their seminal review, Cavanagh and Frank (2014) related theta to cognitive control by showing that theta power increased for novel and difficult stimuli, as well as for negative feedback (stimulus-locked) and errors (response-locked). An emerging consensus suggests that cortical theta coordinates long-range interactions for implementation of cognitive control. For example, using EEG in a Stroop task, Hanslmayr et al. (2008) showed that phase coupling between the anterior cingulate cortex and the left prefrontal cortex lasted longer for incongruent stimuli than for neutral or congruent ones. Several other studies have described theta frequency phase locking between frontal and other areas in humans using EEG (Cohen, 2009; Nigbur et al., 2012; van de Vijver et al., 2011), as well as in non-human animals using intracranial recordings (Narayanan et al., 2013; Phillips et al., 2014). In short, recent literature suggests that theta plays a fundamental role in the exertion of cognitive control.

Alpha band (8-12 Hz) activity has also been implicated in cognitive control. Two main effects are found when considering the interplay between alpha and cognitive control. First, alpha power increases in task-irrelevant brain areas (Klimesch, 1999; Mazaheri et al., 2009; Pfurtscheller, 2003). To explain this increase, Jensen and Mazaheri (2010) proposed that alpha activity blocks irrelevant processing pathways, which in turn allows the relevant brain areas to process the presented information more efficiently. This hypothesis was named ‘gating by inhibition’. Evidence comes from a study by Jokisch and Jensen (2007), in which participants completed a delayed-match-to-sample task, discriminating either the identity or the orientation of a presented picture of a face. Consistent with gating by inhibition, the authors found that alpha activity increased in the brain area that was irrelevant for the current condition. Thus, an increase in alpha power might be linked to inhibition of a targeted (task-irrelevant) area.

Second, alpha power decreases in task-relevant brain areas (Händel et al., 2011; Jensen & Mazaheri, 2010; Thut, 2006). Using a go-no-go task, Mazaheri et al. (2009) observed alpha suppression in task-relevant brain areas. Thus, whereas cognitive control is typically accompanied by high FM theta power, it usually co-occurs with low posterior alpha power.

Generally, both theta and alpha are modulated by cognitive control, albeit in a different direction (in- and decrease for theta and alpha, respectively). Also on a trial-to-trial basis, theta and alpha power are (negatively) correlated. Mazaheri et al. (2009) observed in their go-no-go task that post-trial anterior theta power was negatively correlated with post-trial posterior alpha power, and especially after error trials.

Given these tight couplings between theta and alpha, what is their respective role in cognitive control? One possibility is that theta and alpha simply reflect the anterior and posterior signatures of control measured on the human scalp, respectively. In this view, theta and alpha would always be correlated across time and conditions, yet originate from different neural areas. An alternative view is that theta and alpha both implement control, but at different timescales. This would not be visible with standard paradigms where the involvement of theta and alpha is not tracked across time for the same stimuli. Indeed, the transition from controlled to automatic processing may require many repetitions and consist of several stages (Ashby et al., 2007), with different oscillations potentially involved at different stages. Current protocols cannot disentangle these possibilities.

To address this problem, we devised a paradigm that is able to investigate how cognitive control unfolds over both a fast and a slow timescale. We started from a stimulus-action learning paradigm by Ruge and Wolfensteller (2010). In their fMRI experiment, subjects learned four unique stimulus-action mappings at the start of each experimental block through instruction. Each stimulus-action mapping was shown eight times within one experimental block. The authors investigated changes in neural activity as a function of the number of times a stimulus was seen within a block. On the behavioral level, subjects improved with practice, as indicated by both faster reaction times (RTs) and fewer errors. On the neural level, the authors found that areas that were hypothesized to be involved in automatization (striatum and the pre- and postcentral gyri) became more active with practice. In contrast, brain areas that were presumably related to cognitive control (such as the lateral prefrontal cortex and the intraparietal sulcus) became less active with increasing practice.

In the current work, we investigate the timescales of cognitive control, and its transition into automatic processing, and in particular the role of theta and alpha oscillations in this process. For that purpose we presented one set of four stimulus-action mappings repeatedly within a block (fast time scale, as in Ruge and Wolfensteller, (2010)). The same set of mappings also repeated across blocks (slow time scale), interleaved with blocks where novel mappings were presented. Furthermore, we measured EEG, which allowed us to track theta and alpha power changes over the two timescales, and investigate if theta and alpha show the same pattern across the two timescales, or not.

## Materials and Methods

The experimental procedure was approved by the Ghent University Ethical committee (Faculty of Psychology and Educational Sciences). All 30 participants signed the informed consent. In return for their participation, a monetary compensation of €25 was provided. All participants reported to be free from neurological conditions and had normal or corrected-to-normal vision. All participants were Dutch native speakers, and three of them were left-handed. The experiment started either at 9:00 A.M. or 1:00 P.M. and always lasted two hours. Data from 24 subjects (18 – 47 years old, 23.1 ± 5.6 (mean ± SD) years (17 females, 22 right-handed) were analyzed and reported. Criteria for exclusion were 1) data loss (6.66%), 2) inferior data quality (10%), and 3) technical issues (3.33%). The experiment was created using PsychoPy 3 (Peirce, 2007).

### Design and Experimental Protocol

The experiment took place in a Faraday cage and consisted of a learning task. Subjects learned stimulus-action mappings through trial-and-error learning. Subjects responded by pressing the “f” and “j” keys on the keyboard of the experiment computer using their left and right index fingers respectively. Each experimental block started with four stimuli that were presented vertically (familiarization phase; no action mapping information provided; see Figure 1A). Participants were instructed at the beginning of the experiment that these would be the stimuli that they would encounter in the upcoming block.

**Figure 1.**
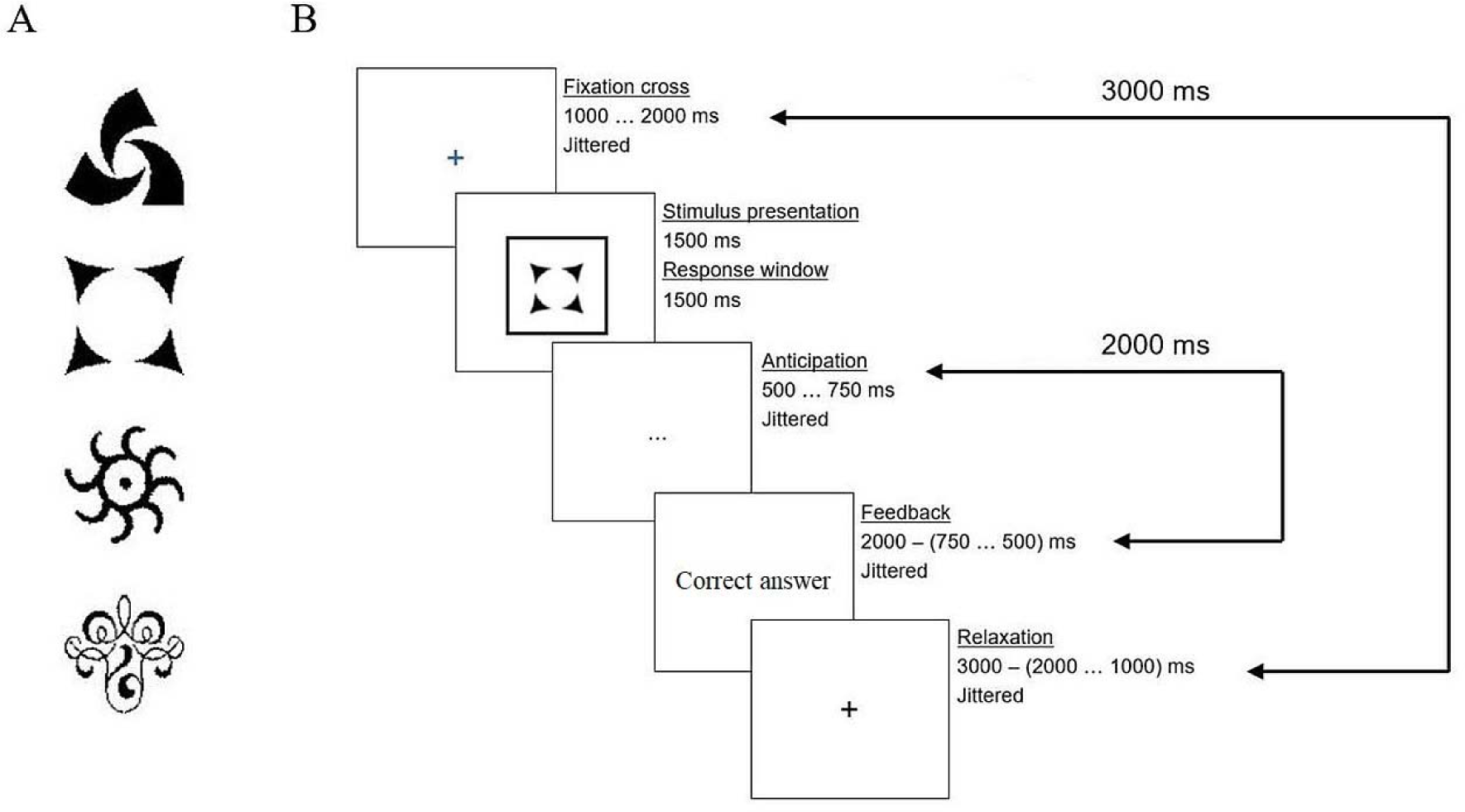
Visualization of the stimuli seen during the experiment, and a trial overview. (A) An example of the stimulus overview that was provided at the start of each experimental block. This overview represented the four stimuli that would be shown in the following block. (B) An overview of an experimental trial. Each trial lasted exactly 6500 ms in total.

After stimulus inspection, participants could start the experimental block by pressing the space bar on their keyboard. Following this, a five-second counter started, and the first trial of that block would start. A trial started with a fixation cross colored in either blue or black, balanced across subjects. The participants were informed prior to the start of the experiment what these colored fixation crosses meant. For half of the subjects, a blue fixation cross meant that a stimulus was about to appear, while a black fixation cross signified the end of the trial. For the other half of subjects, the opposite pattern applied. The duration of the first fixation cross was temporally jittered, meaning that the presentation time was drawn from a uniform distribution (1000-2000 milliseconds (ms)). Then, participants saw the stimulus framed by a black box in the center of the screen. From that moment, the subject had 1500 ms to press one of the two response buttons. Only when a button was pressed in time, the box would turn grey, indicating that an answer was registered. After 1500 ms, the stimulus disappeared from the screen, and was replaced by three dots which we will refer to as the ‘anticipation phase’. This stimulus timing was again temporally jittered, with a presentation time between 500 and 750 ms. Following the anticipation phase, the feedback (right, wrong, or too late) was provided in the form of a simple Dutch text message (“Correct answer”, “Wrong answer”, “Too slow”) in the middle of the screen. The feedback presentation time depended on the presentation time of the anticipation phase. Together, these two phases lasted for 2000 ms. Finally, as mentioned earlier, a colored fixation cross marked the end of the trial. The presentation time of this fixation cross depended on the presentation time of the first fixation cross: Together they were shown for 3000 ms. Figure 1B provides a visual representation of a trial in our experiment. Subjects learned the correct stimulus-action mappings via trial-to-trial feedback.

The stimuli in each block were presented in semi-random order. Specifically, stimulus presentation order was generated so that stimuli that were not presented recently had a higher likelihood to be shown in the next trial. Thus, trial number in an experimental block and the number of times a stimulus is seen in a block are positively correlated. The first two blocks (64 trials in total) were performed as practice before the EEG set-up. Each block was comprised of four stimuli that were each shown eight times. Since one stimulus is shown per trial, this makes up a total of 32 trials per block similar to the paradigm described by Ruge and Wolfensteller (2010). The experiment (without practice) lasted for 16 blocks, summing to 512 trials in total per subject. Note that at the end of this experiment a subject had seen 36 (4 times 9) unique stimuli. Of these 36 stimuli, four stimuli served as repeating stimuli (with constant response mapping), while the other 32 were unique (‘novel’) stimuli. Each repeating stimulus was seen 64 times, while the novel stimuli were seen eight times each. The stimuli used in an experimental run were randomly drawn from a collection of 124 stimuli. Half of the blocks (256 trials) contained repeating stimuli; the other blocks contained novel stimuli. The order in which experimental blocks were presented was again semi-random: participants would never see more than two blocks in a row of the same condition (novel/repeating). Participants could take a break at the end of each block, which was denoted by a message on the screen. No time limits were imposed on these breaks. When they were ready to continue, subjects could press the space button, after which they would see the stimuli from the next block. Participants used a cushioned chinrest during EEG recording.

### Data Recording and Initial Processing

All EEG data were collected using a 64-channel BioSemi Active Two System (BioSemi, Amsterdam, Netherlands) at 1024 Hz. The electrodes were placed following the standard international 10-20 electrode mapping defined by Jasper (1958). The cap contained a posterior CMS-DRL electrode combination. Six additional external electrodes were placed on the left and right mastoid, lateral canthi of both eyes, and above and below the left eye. The data collected using these electrodes were used as a reference during analysis (the average of the mastoid electrodes), or for detecting and correcting eye movements and eye blinks (the other external electrodes). Data preprocessing was done using the MNE Python software (Gramfort et al., 2014), and custom code. Our pipeline was inspired by the EEGLAB preprocessing pipeline (Delorme & Makeig, 2004), but some adaptations were made. First, the data were re-referenced to the average of the mastoid electrodes. Data was then visually inspected to mark bad channels, which were then interpolated. For 66.66% of analyzed subjects at least one channel was interpolated, with a maximum of six interpolated channels across subjects. The majority of interpolated electrodes were located in the posterior area of the scalp. The next step was to high-pass filter this data at 0.1 Hz, and to split this filtered data in epochs locked to stimulus onset. We then performed a visual inspection of these epochs and excluded the noisy epochs from further analysis. This exclusion process was performed blind towards the experimental conditions. Our next step would be to run ICA on the data epochs. However, Winkler et al. (2015) demonstrated that the efficiency of ICA in reducing artifacts greatly depends on the chosen preprocessing procedure. Specifically, the authors suggest that high-pass filtering the data above 1 or 2 Hz prior to ICA consistently leads to good results. Thus in order to improve our artifact reduction procedure, our next step was to go back to the continuous EEG data obtained after bad channel interpolation. We then bandpass filtered this data between 1 and 40 Hz, and we again split this filtered data in epochs locked to stimulus onset. We then removed the epochs that were marked as bad when visually cleaning the 0.1 Hz high-pass filtered epochs. Following this we used the extended Infomax independent component analysis (ICA) algorithm implemented in MNE (Lee et al., 1999) on the bandpass-filtered, epoched data. These ICA components were then used on the 0.1 Hz high-pass filtered data to remove eye movement artifacts. The final step in our preprocessing pipeline was to remove epochs based on the behavioral data: we removed 1) epochs where the RT deviated more than three standard deviations from the overall mean RT, and 2) epochs where the participants were incorrect, even though they saw that stimuli 3 times or more (assuming those represented attentional slips). The same pipeline was used to preprocess epochs locked to the response. Overall, the complete data cleaning procedure led to an exclusion ranging from 3.32% to 29.30% (*M* = 17.34, *SD* = 5.21) of the recorded epochs across participants.

### Statistical Analysis

#### Behavioral Analysis

All statistical analyses of the behavioral data were performed using the ‘lme4’ package (Bates et al., 2007), which is embedded in the R software (Version 3.6.3). All associated plots were created using the visualization libraries ‘seaborn’ and ‘Matplotlib’ (Hunter, 2007) in the Python programming language (Version 3.5.6).

For the behavioral results, two dependent variables are considered: RT and error rates. RT was first log scaled prior to statistical tests to increase normality. For each statistical test involving RT as the dependent variable, we built a linear mixed effects model. For errors, a generalized linear mixed effects model was utilized. All models contained a random intercept for subject. The independent variables were always included as fixed effects. Note that in plots the original, untransformed, data is displayed.

In order to elucidate the effects over the fast timescale, we analyzed how the recorded behavioral and neural metrics evolved within blocks. To this end, we analyzed dynamics as a function of stimulus number, i.e. the number of times a stimulus was seen by a subject. For example, when investigating how RT evolved over the fast timescale, we would build a linear mixed effects model, where the independent variables were stimulus number and condition, and the dependent variable RT.

The slow timescale represents changes between blocks. To capture this, we analyzed changes in our recorded data as a function of block number and condition.

#### EEG Data Analysis

Our analyses focus on the total power, but we also investigate evoked power. Evoked power refers to the phase-consistent part of the EEG signal (Hajihosseini & Holroyd, 2013), which can be accessed by computing the event-related potential (ERP) before spectral analysis.

Before exploring how alpha and theta power fluctuate across timescales, additional processing was done. The nature of this data preparation depended on several factors. First, the timescale (fast or slow) considered in the specific analysis influenced our data analysis. Second, whether we looked at either total power or evoked power impacted our analysis choices. The third and final factor of our pipeline was the event (stimulus or response) to which the data was time-locked. We will now delineate how each preprocessing decision was implemented in our analyses.

To investigate how oscillatory alpha and theta power evolve for each timescale, we again used stimulus numbers and block numbers as independent variables for the fast and slow timescales, respectively. Specifically, we started our analysis of oscillatory power by comparing the power values for the end points on each timescale. For the fast timescale, we compare the recorded spectral power for stimulus 1 versus 8. For the slow timescale, the power in experimental block 1 is contrasted with the power in block 8. Thus, in order to equate the relevant power measures, we first split the recorded EEG data for each subject in two extreme parts for each timescale: data representing earlier timepoints (stimulus 1, or block 1) and data from late timepoints (stimulus 8, or block 8). Then, we computed the time-frequency representation (TFR) for each subset of the data on the same timescale using Morlet wavelets. The power estimates were computed for 15 frequency bands, log spaced between 4 Hz and 30 Hz. The number of cycles used for each frequency was equal to the frequency divided by two, to preserve the balance between temporal and frequency precision (Cohen, 2014). This procedure ultimately resulted in TFRs for each timepoint for each subject. To compare the resulting TFRs, a cluster-level statistical permutation test was used to account for multiple comparisons (Maris & Oostenveld, 2007). This permutation test was run with 1000 permutations, and a significance threshold of 5% was used.

For evoked power, the aforementioned permutation test was run on the TFRs of the ERPs of specific subsets. For the fast timescale this meant that we only included the data representing stimulus 1 and 8 in the permutation test. For each of these subsets the ERP was computed, which was then followed by initializing the permutation test. A similar procedure was followed for the data tracking the slow timescale, but there the subject-specific data was split in a part containing data of block 1, and a part containing the data of block 8. With respect to total power, we followed the same procedure, except for omitting the ERP computation step.

Stimulus-locked epochs were 2500 ms long (−1000 to 1500 ms relative to stimulus onset). We analyzed the time period ranging from 0 to 1000 ms after stimulus onset for the fast timescale, and the period from 0 to 750 ms after stimulus onset for the slow timescale. The period from 1000 to 250 ms before stimulus onset served as the baseline. For response-locked data, we also used a pre-stimulus baseline period, but this one ranged from 1000 to 500 ms before stimulus onset. We shortened our baseline period because our analyzed time period was shorter: we only consider the data from 235 ms before response onset, to 305 ms after response. The response-locked time window was defined like this because otherwise some overlap might occur with the anticipation phase of the next trial (see Figure 1B), especially for the trials where subjects exhibited later responses.

Based on the results of the cluster-level statistical permutation test, time-frequency plots were created. The time-frequency plots represent the cluster statistics averaged over all channels. This resulted in a two-dimensional plot displaying the computed statistics as a function of time and frequency bands. To assess where power fluctuations originate from on the scalp, we also created topographical plots by subtracting block 1 (stimulus 1) from block 8 (stimulus 8).

In summary, depending on which effects we investigated changes in the processing pipeline were implemented. When the timescale of interest (fast or slow) was changed, this impacted which TFRs were compared using the cluster-level statistical permutation test. Additionally, investigating fluctuations in either total or evoked power impacted on which data the aforementioned TFRs were computed. Finally, baseline correction was always applied but the time window used depended on whether the data was stimulus-locked or response-locked.

In the time-frequency clustering algorithm, we only contrasted stimulus 1 or block 1 with stimulus 8 or block 8 respectively across both timescales. However, we conducted follow-up analyses on the intermediate stimuli 2-7 and blocks 2-7 (thus avoiding “double dipping”; Kriegeskorte et al., 2009). To this end, we used the ‘lme4’ package (Bates et al., 2007), incorporated in the R software (Version 3.6.3). For example, consider the hypothetical situation where a significant difference in theta power is found in some time window after stimulus onset in the stimulus-locked cluster test comparing stimuli 1 and 8. The first step in the follow-up analysis would be to compute the average theta power in that same time window across trials. Next, we would exclude all the trials that contain data for stimuli 1 and 8. Finally, we would build a linear mixed effects model with theta power as dependent variable, subject number as a random intercept, and stimulus number and condition as fixed effects. A similar procedure was used for analyzing effects over blocks 2 to 7.

## Results

### Behavioral Results

Focusing on the fast timescale, stimulus number had a main effect on both RT, *F*(7, 5561) = 61.48, *p* < .001, and error rate, *X^2^*(7, N = 24) = 122.38, *p* < .001. Both RT and error rate decreased as stimulus number increased. How RT and error rate evolved over the fast timescale is visualized in Figure 2A and 2B, respectively. Note that accuracy starts off slightly below 50% because of a dependency between stimuli: If the participant has already seen three out of four stimuli (e.g., two with left and one with right response), the response to the fourth stimulus can be known with certainty (in this example, right response).

**Figure 2.**
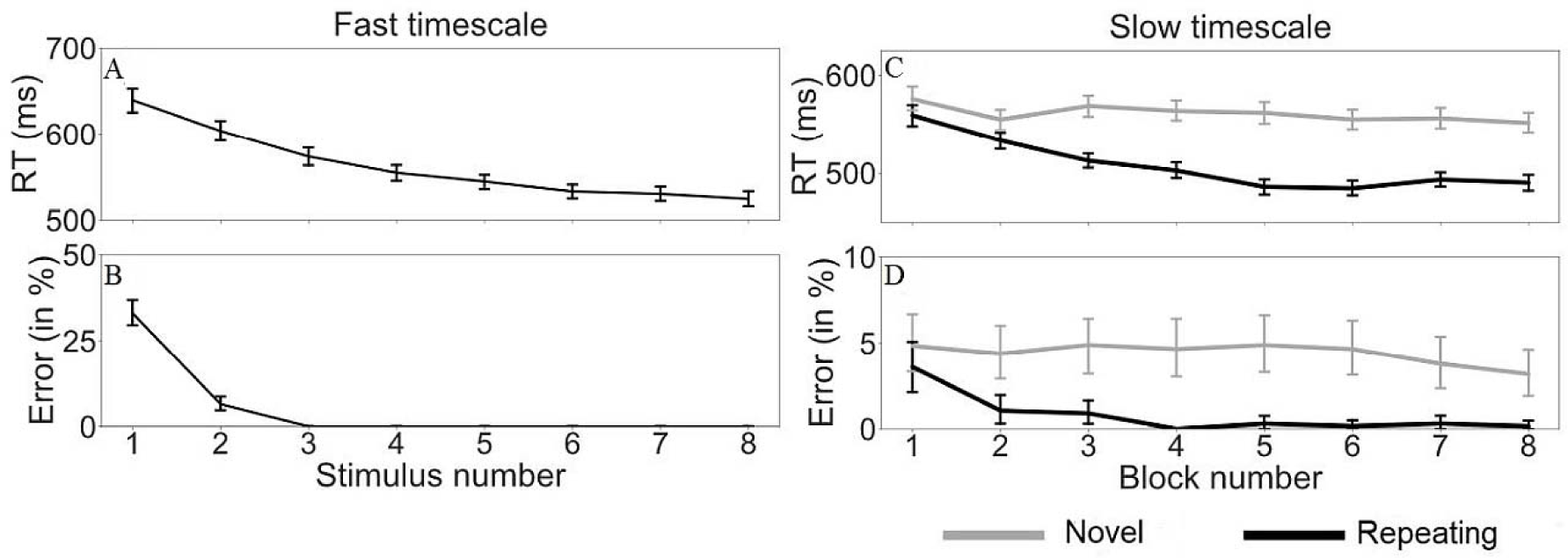
RT and error rate following the fast and the slow timescale. RT (A) and error rate (B) as a function of stimulus number, respectively. These plots capture behavioral effects following the fast timescale. RT (C) and error rate (D) as a function of block number, respectively. Thus, these plots show behavioral changes on the slow timescale. A distinction is made based on condition, where ‘novel’ indicates that new stimuli were shown in that block, and ‘repeating’ means that stimuli were shown that returned throughout the experiment. The vertical lines represent bootstrapped 95% confidence intervals in all plots.

Next, we considered the same dependent variables on the slow timescale. Both block number, *F*(7, 10118) = 25.79, *p* < .001, and condition (novel or repeating), *F*(1, 10118) = 542.87, *p* < .001, had a significant impact on RT. Additionally, the interaction effect between block number and condition also proved significant, *F*(7, 10118) = 14.50, *p* < .001. A follow-up pairwise t-test (Benjamini-Hochberg adjusted) showed that in the repeating condition, block 1 was significantly different from all others (all *p* < .001). In the novel condition, none of the blocks differed significantly (all *p* > .05). On error rate, block number had a significant impact, *X^2^*(7, N = 24) = 34.29, *p* < .001. The main effect of condition on error rate proved to be non-significant, *X^2^*(1, N = 24) = 0.03, *p* = .871. However, the interaction effect of block number and condition was significant, *X^2^*(7, N = 24) = 27.30, *p* < .001. Specifically, using the same follow-up test, we concluded that in the repeating condition the error rate in block 1 was significantly higher than the error rate in the other blocks (all *p* < .001), but remained constant in the novel condition (all *p* > .05). How RT and error rate evolved following the slow timescale is shown in Figure 2C and 2D, respectively.

### EEG Results

#### Fast timescale

In total power changes on the fast timescale (stimulus 1 vs. stimulus 8), we found a significant cluster in the stimulus-locked data. Specifically, we noted a significant cluster after cluster correction (*p* = .001) encompassing both the alpha band (8 – 12 Hz) and part of the beta band (14 – 30 Hz) approximately between 700 and 850 ms after stimulus onset (Figure 3A). Since two separate frequency bands appear to be involved, we conducted separate analyses for each band. The remainder of this paragraph is focused on the alpha band, after which we turn to changes in beta power. The alpha modulation is localized in the posterior part of the scalp (Figure 3B). Follow-up analyses focusing on alpha power (stimulus 2 to 7) in this significant cluster, indicated that alpha power (from here on abbreviated as alpha) significantly increased with stimulus number, *F*(5, 7684) = 20.34, *p* < .001. The main effect of condition also proved to be significant, *F*(1, 7684) = 61.70, *p < .*001. Finally, the interaction effect between stimulus number and condition was also significant, *F*(5, 10118) = 3.40, *p* = .005. How alpha evolves as a function of stimulus number and condition is visualized in Figure 3C. An additional analysis was carried out to investigate how alpha behaved at the single-electrode level. To this end we analyzed alpha as measured on electrode Pz (based on Figure 3B). Since the single-electrode results were similar to the ones visualized in Figure 3C, we refer the interested reader to panels A and B of Figure S1 in the Supplementary Materials section. Finally, we note that if we restrict our analyses to only the electrodes that made a significant contribution we obtained highly similar results. The comparison between the results based on all 64 electrodes and the results obtained when only taking into account the significant electrodes is visualized in Panels A and B respectively of Figure S2.

**Figure 3.**
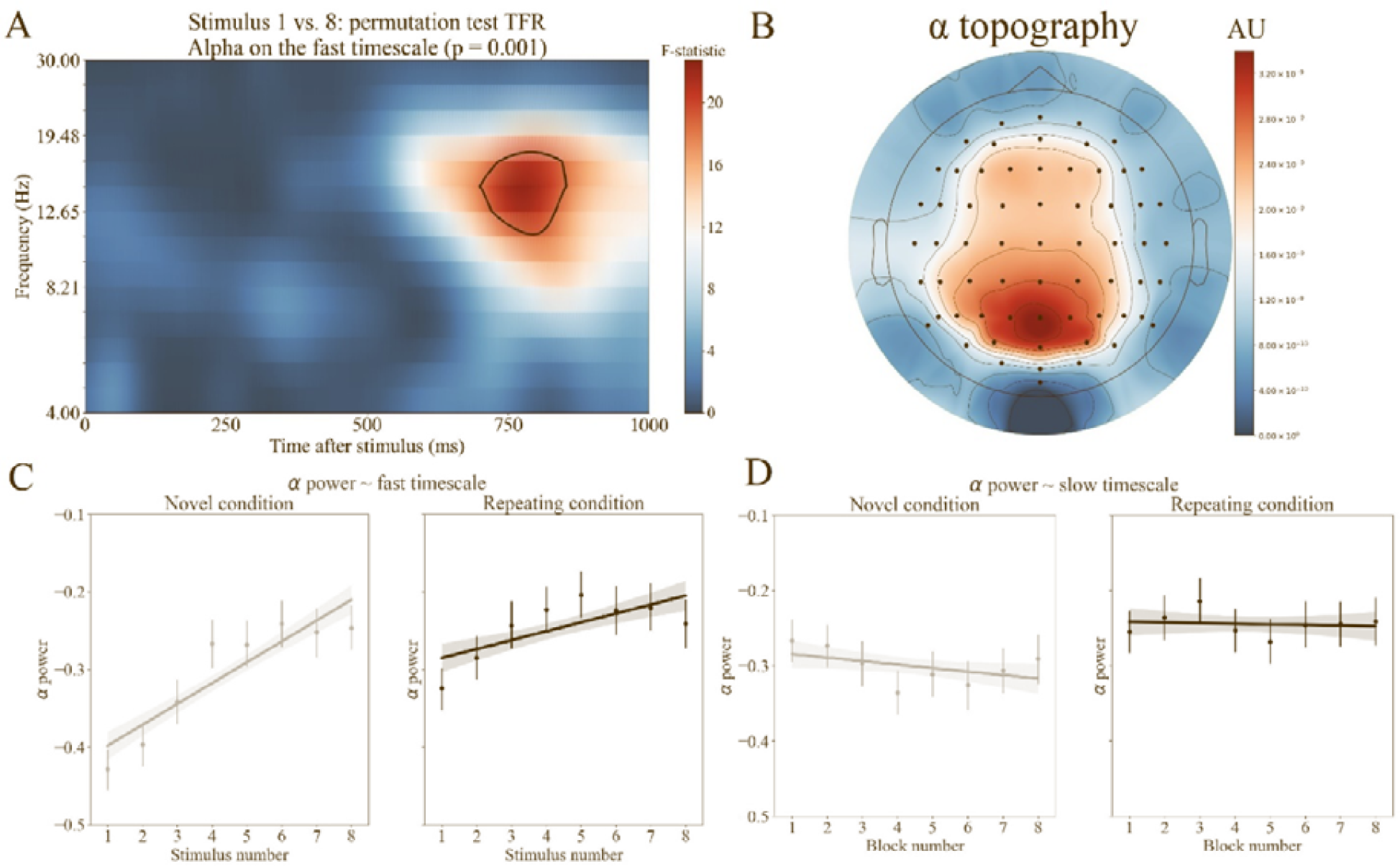
Alpha power across timescales. (A) This time-frequency plot depicts the cluster statistics which were averaged over all channels, resulting in a two-dimensional plot displaying the computed statistics as a function of time and frequency band. We delineated the 95% largest F-values. Note that the delineated area only serves a visualization purpose, and is not used in any follow-up analyses described later. (B) This plot depicts the topography of the difference of stimulus 8 minus stimulus 1 in the alpha frequency band, averaged across time between 700 and 850 ms after stimulus onset. (C) Alpha (8 – 12 Hz) between 700 and 850 ms after stimulus onset averaged across time and trials as a function of stimulus number. The analyzed time-interval was defined based on panel A. (D) Alpha averaged across time and electrodes between 250 and 400 ms after stimulus onset as a function of block number and condition. We analyzed this time period based on the results depicted in Figure 4A. Note that in both panels C and D averages are represented by dots, while bootstrapped 95% confidence intervals are represented by vertical lines. The straight lines represent linear regression lines.

Following this, we investigated potential shifts in beta power (from here on referred to as beta). This follow-up analysis pointed that beta significantly increased as a function of stimulus number, *F*(5, 7684) = 25.91, *p < .*001. Also the main effect of condition, *F*(1, 7685) = 69.55, *p < .*001, and the interaction effect between stimulus number and condition, *F*(5, 7684) = 3.63, *p* = .003, proved to be significant.

Next, we analyzed stimulus-locked evoked power contrasting stimulus 1 vs. 8. We found a significant alpha cluster between 700 and 800 ms after stimulus onset (*p < .*001). Combined with the previous result, this indicates that the stimulus-locked alpha burst can be seen in both the total and the evoked power, occurring more than 700 ms after the stimulus onset.

For the response-locked data, we again started by comparing the total power measures for stimulus 1 vs. stimulus 8, performing the same cluster-correction procedure. A significant alpha cluster was observed (*p* = .002) around 200 ms after response onset. A visualization of this response-locked alpha cluster can be seen in Figure S3. No significant results were obtained when analyzing the response-locked, evoked power.

#### Slow timescale

For the slow timescale, we started by analyzing the total power fluctuations by contrasting block 1 with block 8 (block 1 vs. block 8). The permutation test highlighted two significant theta clusters (4 – 8 Hz), one at stimulus onset and one around 400 ms after stimulus onset (*p* = .04) (Figure 4A). The dynamics of the cluster in the time window from 250 to 450 ms originate from the frontal scalp region (Figure 4B). A follow-up test (blocks 2-7) using a linear mixed effects model showed that block number is a significant predictor of theta power, *F*(5, 7651) = 2.59, *p* = .02. In contrast, the main effect of condition was not significant, *F*(1, 7561) = 1.12, *p* = .29. The interaction effect between condition and block number, *F*(5, 7651) = 2.32, *p* = .04, proved significant (Figure 4D). Similar to the fast timescale, an additional analysis was carried out to elucidate how theta behaved at the single-electrode level (electrode Fz, chosen based on Figure 4B). Since the single-electrode results were similar to the ones visualized in Figure 4D, we refer the interested reader to panels C and D of Figure S1. Finally, in line with the fast timescale, we compared the results obtained based on the full electrode set with the outcome based on the significant electrodes. This again yielded nearly identical results. The difference for results obtained from the full set vs. the significant subset can be seen when contrasting panels C and D of Figure S2 respectively.

**Figure 4.**
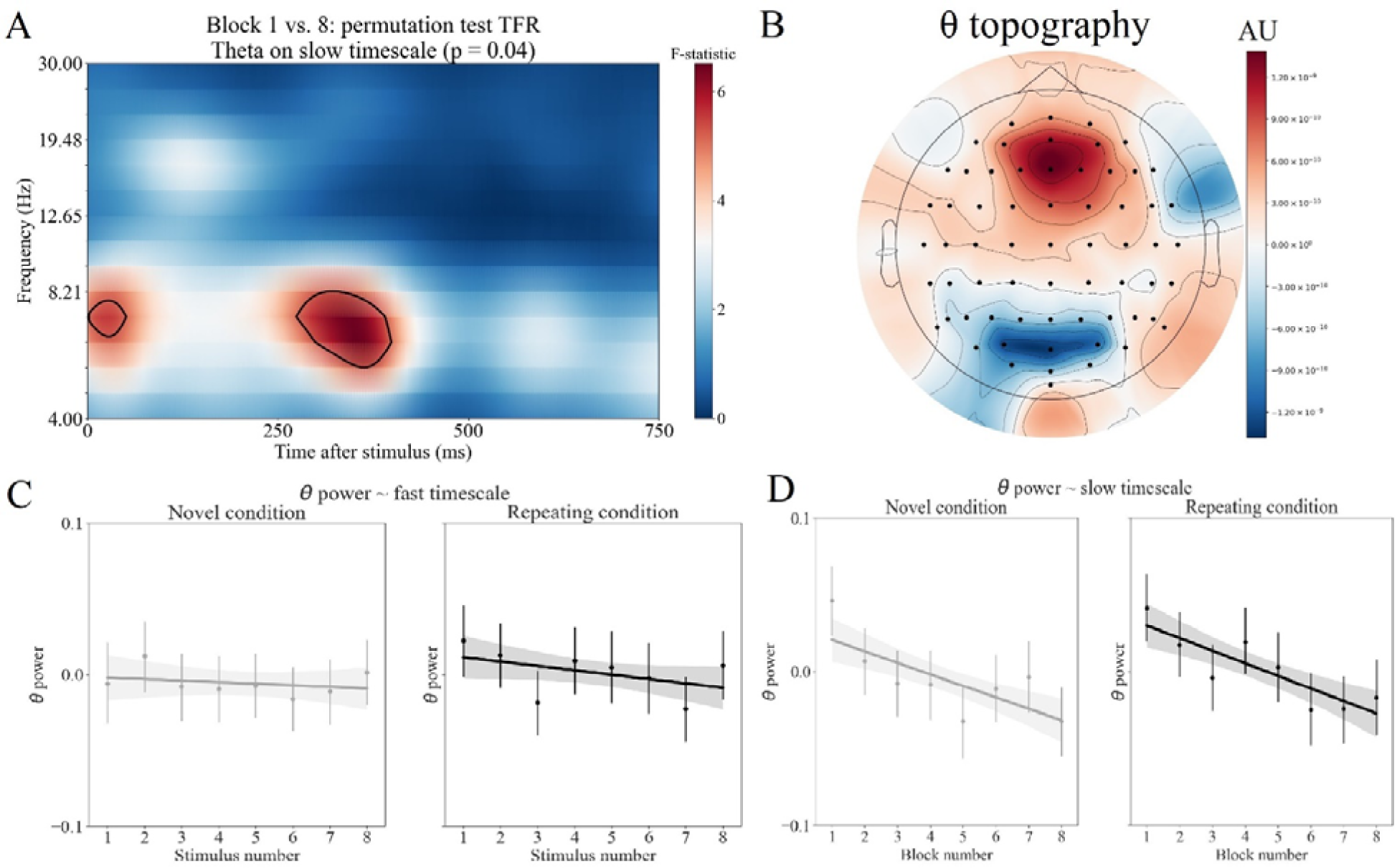
Theta power across timescales. (A) Channel-averaged time-frequency representation of power. Black curves indicate statistically significant clusters. We delineated the 95% largest F-values. Note that the delineated area only serves a visualization purpose, and is not used in any follow-up analyses described later. (B) This figure depicts the topography of the contrast block 8 minus block 1 in the theta frequency band, averaged across time in the period between 250 and 400 ms after stimulus onset. We specifically selected this time window and frequency band based on the results of the permutation test depicted in panel A of this figure. We highlight an anterior positive difference and a posterior negative difference. (C) Theta (4 – 8 Hz) between 700 and 850 ms after stimulus onset averaged across time and subjects as a function of stimulus number. We analyzed this time period based on the results depicted in Figure 3A. (D) Theta averaged across time and electrodes between 250 and 400 ms after stimulus onset as a function of block number and condition. The analyzed time-interval was defined based on panel A. Note that in both panels C and D averages are represented by dots, while bootstrapped 95% confidence intervals are represented by vertical lines. The straight lines represent linear regression lines.

We also analyzed the contrast between block 1 and block 8 with a focus on stimulus-locked evoked power, response-locked total power and response-locked evoked power. None of these contrasts yielded significant results.

#### Alpha-theta Dissociation

In order to investigate whether the fluctuations in alpha and theta power are specific to the fast- and slow timescales respectively, we additionally tested the interaction between stimulus (block) number and frequency band on each time scale separately. On the fast timescale, the interaction between stimulus number and frequency band (theta, alpha) indeed proved significant, *F*(7, 20259) = 22.33, *p < .*001. Similarly, for the slow timescale the interaction effect between block and frequency band (theta, alpha) also proved to be significant, *F*(7, 20259) = 2.88, *p* = .005. The current interactions, however, may originate from the fact that each time point (stimulus repetition or block) was coded as a separate factor in the analysis. Thus, to further clarify these results, we computed the subject-specific slope of the least-squares regression lines created when regressing alpha (or theta) power as a function of stimulus (or block) number. Then, we compared the subject-specific slopes for the novel condition with the slopes for the repeating condition using a paired samples t-test. This test was performed for each combination between frequency band and timescale. We found only one significant difference in subject-specific slopes between conditions: when considering alpha as a function of stimulus number we found that the subject-specific slopes for the novel condition (*M* = 0.03, *SD* = 0.02) compared to the repeating condition (*M* = 0.01, *SD* = 0.02) differed significantly, *t*(24) = 3.24, *p* = .004. All other post-hoc tests yielded non-significant results (all *p* > .1). The results of this post-hoc analysis confirm that only the interaction effect between stimulus number and condition on alpha survives after further statistical testing.

As another follow-up test we investigated whether alpha (8 – 12 Hz) significantly changed over the slow timescale. To investigate this, we considered the time period from 700 to 850 ms after stimulus onset, since significant alpha effects were found in this window on the fast timescale. This analysis indicated that alpha did not change as a function of block number, *F*(7, 10118) = 1.57, *p* = .14. The main effect of condition proved significant, *F*(1, 10118) = 69.06, *p < .*001. Finally, the interaction between block number and condition was not significant, *F*(7, 10118) = 1.73, *p* = .10. How alpha behaved on the slow timescale is depicted in Figure 3D. Again, in order to explore potential power fluctuations in the beta band, we conducted an exploratory analysis to investigate how beta changes on the slow timescale. This analysis showed that block number had no significant main effect on beta, *F*(7, 10119) = 1.45, *p* = .18. The main effect of condition reached statistical significance, *F*(1, 10119) = 79.14, *p* < .001. The interaction effect between block number and condition did not prove to be significant, *F*(7, 10118) = 1.88, *p* = .07. This analysis suggests that alpha and beta behave in a similar fashion on the slow timescale. Next, we investigated whether theta frequency effects (4 – 8 Hz) could be observed on the fast timescale. To this end, we analyzed theta power in the time window ranging from 250 to 400 ms after stimulus onset. This time window was chosen based on our analyses for the slow timescale, which are described in the previous section. We found a significant main effect of stimulus number on theta power, *F*(7, 10118) = 2.04, *p* = .05 (Figure 4C). Both the main effect of condition, *F*(1, 10118) = 1.12, *p* = .29, and the interaction between stimulus number and condition, *F*(7, 10118) = 0.89, *p* = .51, were not significant. Thus, we argue that on the fast time scale, only alpha changed; whereas on the slow time scale, only theta changed across time.

#### Correlation between neural- and behavioral metrics

In a final analysis step, we investigated whether alpha and theta can act as predictors for RT. To explore this, we created a linear mixed effects model with RT as dependent variable. This model had only one random intercept, namely a subject identifier. Both theta and alpha, denoting power in a specific time window as described earlier, were included as random slopes. Furthermore, the model counted five fixed effects: theta, alpha, stimulus number, block number and condition. Finally, two interaction effects were added: the interaction between stimulus number and condition and between block number and condition. Our main goal of this analysis was to investigate whether alpha and theta had a significant contribution in explaining RT. After fitting this model, we performed a model selection procedure using the AIC as criterion. The resulting model only contained theta, suggesting that theta, but not alpha, is a significant predictor of RT, *F*(1, 22) = 9.08, *p* = .007.

## Discussion

Previous research indicates that alpha and theta frequencies play instrumental roles in cognitive control, but it remains unclear how they unfold during the implementation of cognitive control. In this study, participants learned a set of stimulus-action associations. These associations were block-specific and could be either novel or repeating. Learning took place on two separate timescales: a fast (within blocks) timescale, and a slow (between blocks) timescale. Our results indicate that cognitive control is implemented across both a fast and a slow timescale. On the fast timescale we observed that alpha power significantly increases over time, and more strongly so in the novel than in the repeating condition (see Figure 3C). Interestingly, beta behaved similarly: It also increased over time with a stronger increase in the novel condition compared to the repeating condition. A significant alpha cluster was also found in the response-locked data, indicating that the stimulus-locked effects could not be explained by shifts in RT as the experiment progresses. In contrast, on the slow timescale theta power decreases over time (see Figure 4D), but not differently across conditions. Note that in this contrast the interaction effect between condition and time (block number in this case) proved significant. However, since this effect is comparatively small (p = .04) and since the plot (Figure 4D) gives no visual indication of an ongoing interaction effect we refrain from interpreting this effect albeit its statistical significance. We await more data to check for robustness of this interaction effect. Alpha did not significantly change in the slow timescale, whereas theta significantly decreases on the slow timescale. Follow-up tests show significant interactions between stimulus number and frequency band on the fast time scale, showing that alpha increased faster than theta on the fast time scale; and between block and frequency band on the slow timescale; showing that theta decreased faster than alpha on the slow time scale. Taken together, we argue that alpha increases are specific to the fast timescale, whereas a decrease in theta power is specific to the slow timescale.

We interpret the results obtained at the fast (within a block) timescale in the ‘gating by inhibition’ framework proposed by Jensen and Mazaheri (2010), according to which alpha oscillations suppress task-irrelevant brain areas, and thereby indirectly ‘gate’ task-relevant brain areas. Evidence for this hypothesis comes from studies in memory (Grabner et al., 2003; Sauseng et al., 2005), face processing (Jokisch & Jensen, 2007), visual detection (Hanslmayr et al., 2005), and cognitive control dynamics (Carp & Compton, 2009; Compton et al., 2011; Janssens et al., 2018). For example, Carp and Compton (2009) and Compton et al. (2011) observed a significantly lower alpha power in the inter-trial interval of a Stroop task in error relative to correct trials. Similar results were obtained when contrasting high-conflict with low-conflict trials. From the ‘gating by inhibition’-framework, alpha may reflect inhibition of incorrect stimulus-action mappings.

The alpha effect occurred in the time window 700 to 850 ms after stimulus onset, well after response (median RT was 510.06 ms). So if alpha appears after responding, what is it functional role? We hypothetically propose that alpha relates to “neural replay”, defined as structured reoccurrences of spontaneous brain activation patterns specific to recent items, for the purpose of learning (Peyrache et al., 2009). Recently, Higgins et al. (2021) suggest that brain activation patterns associated with replay corresponded with activation of two resting state networks: the Default Mode Network (DMN) and the parietal alpha network. Since several studies (Foxe & Snyder, 2011; Jensen & Mazaheri, 2010; Mazaheri et al., 2009) suggest that alpha has an inhibitory role, and Higgins et al. found that spontaneous replay is associated with heightened activation in parietal alpha, post-response alpha activity might reflect the inhibition of incoming sensory information required to allow for spontaneous replay (and thus learning) of earlier acquired information. This interpretation is consistent with the fact that alpha power increases during a block (more time off and thus more replay possible as time proceeds), and specifically so in the novel condition (when learning and thus replay is needed).

Note that the increase in alpha might reflect an increasingly fast response process as the block unfolds. However, this interpretation is unlikely. Specifically, RT decreases as a function of stimulus number from approximately 700 ms (stimulus 1) to approximately 500 ms (stimulus 8; see Figure 2A). However, the alpha effects are localized in a later time window situated around 800 ms (see Figure 3A), suggesting that these alpha effects occur well after response onset.

One might argue that our results are not in line with the ‘gating by inhibition’-framework (Jensen & Mazaheri, 2010) since this theory proposed a principal role for alpha oscillations, whereas our results point to modulations both in the alpha and beta band. However, the frequency bands observed by earlier work (Jensen & Mazaheri, 2010; Mazaheri et al., 2009) extend well beyond the conventional 12 Hz alpha band, similar to the results presented in this paper (e.g. see Figure 2 panel A of Mazaheri et al.). Hence, our results seem well in accordance with the gating by inhibition account and with earlier data that underlies this account.

We explain our slow (i.e., across blocks) time scale results in the theta band as reflecting the role of FM theta in the exertion of cognitive control. In their review paper, Cavanagh and Frank (2014) observed that cognitive control tasks typically showed a neural signature in the theta band approximately 250 to 500 ms after stimulus onset. We hypothesized that this theta band activation originates from the mid-frontal part of the scalp. In these results, FM theta is interpreted as a signal that marks the need for cognitive control. Other studies focused on the role that FM theta might play in the coordination of cognitive control. For instance, it is thought that (FM) theta synchronizes posterior processing areas depending on the task at hand (Fries, 2015). Several authors (Fries, 2015; Verbeke & Verguts, 2019; Voloh et al., 2015; Womelsdorf et al., 2006) suggested that communication between neural areas is achieved using gamma-band synchronization, and the role of theta is to reset gamma such that the appropriate task-relevant areas may subsequently phase-lock in the gamma band. Theta-band phase synchrony can be observed between central areas and more distant brain areas. For example, after errors, enhanced synchrony is observed between the MFC and the occipital cortex (Cohen, 2009); and after incongruent stimuli during a Stroop task (Hanslmayr et al., 2008), synchrony is increased between FM and frontolateral regions. Consistently, we observed decreased theta power when comparing block 8 minus block 1, in both the novel and the repeating condition. Note that one could hypothesize that these theta effects are response-related. However, this is also unlikely because the theta effects occur well before the mean RT. Thus, since theta decreases equally in both novel and repeating conditions over a slow timescale, our results suggest that theta is implicated in a general cognitive process, such as mapping stimuli on available actions. We hypothesize that the decrease in theta reflects that this mapping process enfolds more efficiently when a subject is more experienced with the task, irrespective of the stimuli or actions that are currently being mapped. In short, theta reflects a general (i.e., not stimulus- or action-specific) learning process that takes place on a slow timescale, and becomes optimized with more training. Finally, the correlation of theta with RT suggests that trials where this mapping process is implemented more strongly, allow more efficient processing and thus faster RT.

Note that both the alpha and the theta effects were found in a restricted time window after stimulus onset. Hence, our results might not be generalizable to time windows different to the ones we discussed here. Future research endeavors might help in determining whether the alpha and theta modulations we found after stimulus onset can also be observed in different time windows.

An alternative interpretation of our results is that they reflect cognitive fatigue resulting from prolonged testing on a simple task. A study by Wang et al. (2016) suggests that this interpretation is improbable. In this EEG study, participants were asked to perform a Stroop task for a duration of 160 minutes with the goal to elicit cognitive exhaustion. Wang et al. (2016) report an anterior ERP that seems to be related to compensatory mechanisms used to counteract the effects of cognitive fatigue. Specifically, the authors note an inverted U-shaped relationship between the time on task and the amplitude of this frontal ERP. The signal peaks when the subject attempts to compensate cognitive fatigue, and decreases both during baseline, and when the subject is too tired to engage in compensation. Crucially, the latter was observed in the time window ranging from 120 minutes to 160 minutes after the start of the experiment. Thus, it seems unlikely that our results can be explained by cognitive fatigue, since the neural markers of fatigue occur after a time-on-task greatly exceeding our time-on-task of 60 minutes.

Our results can also be linked to the literature on automatization and extensive training. For instance, in the fMRI study by Ruge and Wolfensteller (2010) on instruction-based learning, the authors noted that brain activation in the central gyri and the striatum increased on a fast timescale, suggesting that these areas are relevant for automatization. On the other hand, activation in the lateral PFC and the intraparietal sulcus (IPS) decreases with more experience with a stimulus. Given the posterior location of IPS, the fact that increased alpha correlates with decreased BOLD (Ritter et al., 2009), and our hypothesized inhibitory role of alpha, as well as our finding that alpha increases with stimulus experience, we hypothesize that the measured increase in posterior alpha may signify the active inhibition of the IPS. Since the spatial resolution of EEG is too low to verify this theory, a future study might investigate this further using a method that has both high spatial and high temporal resolution (such as intracranial EEG). We furthermore show that theta power changes on a slower timescale, suggesting that automatization might occur on distinct timescales signified by distinct neural markers.

Future research will have to extend our paradigm to one where participants are trained over consecutive days, thus to investigate the neural fate of alpha and theta in extended automatization contexts. Future research might also extend our research findings using source-localization to narrow down the neural sources of our reported alpha and theta activations. These additional manipulations would provide extra information on alpha and theta dynamics, such as at which point in time the power measures stabilize, which deep brain structures are involved, and whether learning and automatization enfold on other timescales than the ones reported in the current study.

## Conclusion

To sum up, we found that theta and alpha frequency both play a role during stimulus-action learning, but their impact differs across timescales. Alpha band dynamics are specific to the fast timescale, whereas theta changes are observed in the slow timescale, and similar for novel and repeating stimuli. Our data suggest that automatization is not a unitary process but consists of at least two dissociable steps. Which neural structures are involved, how these neural processes evolve with more training, and the nature of their computational role, remain to be investigated in future research.

## Acknowledgements

The authors declare no conflict of interest. This work was supported by the EOS grant number G0F3818N by Research Foundation Flanders (to TV, CNB and PH). PV was supported by grant 1102519N from Research Foundation Flanders.

## Author contributions

PH, CNB and TV designed the study. Data was collected by PH and PV, and processed by PH. PH performed the data analyses. PH, CNB and TV interpreted the data. PH and TV wrote the manuscript, incorporating input from all authors. CNB and TV provided supervision during this research project. All authors read and approved the final version of this manuscript.

## Data and code availability statement

The data that support the findings of this study are openly available in Zenodo at 10.5281/zenodo.4659714. All scripts used to preprocess, analyze and visualize our data are stored on GitHub: https://github.com/phuycke/alpha_theta_timescales.

## References

Anguera, J. A., Boccanfuso, J., Rintoul, J. L., Al-Hashimi, O., Faraji, F., Janowich, J., Kong, E., Larraburo, Y., Rolle, C., Johnston, E., & Gazzaley, A. (2013). Video game training enhances cognitive control in older adults. Nature, 501(7465), 97–101. https://doi.org/10.1038/nature12486

Ashby, F. G., Ennis, J. M., & Spiering, B. J. (2007). A neurobiological theory of automaticity in perceptual categorization. Psychological Review, 114(3), 632–656. https://doi.org/10.1037/0033-295X.114.3.632

Bates, D., Sarkar, D., Bates, M. D., & Matrix, L. (2007). The lme4 package. R Package Version, 2(1), 74.

Botvinick, M. M., Braver, T. S., Barch, D. M., Carter, C. S., & Cohen, J. D. (2001). Conflict monitoring and cognitive control. Psychological Review, 108(3), 624–652. https://doi.org/10.1037/0033-295X.108.3.624

Buzsáki, G. (2002). Theta Oscillations in the Hippocampus. Neuron, 33(3), 325–340. https://doi.org/10.1016/S0896-6273(02)00586-X

Carp, J., & Compton, R. J. (2009). Alpha power is influenced by performance errors. Psychophysiology, 46(2), 336–343. https://doi.org/10.1111/j.1469-8986.2008.00773.x

Cavanagh, J. F., & Frank, M. J. (2014). Frontal theta as a mechanism for cognitive control. Trends in Cognitive Sciences, 18(8), 414–421. https://doi.org/10.1016/j.tics.2014.04.012

Cohen, M. X. (2009). Unconscious errors enhance prefrontal-occipital oscillatory synchrony. Frontiers in Human Neuroscience, 3. https://doi.org/10.3389/neuro.09.054.2009

Cohen, M. X. (2014). Analyzing neural time series data: Theory and practice. MIT press.

Compton, R. J., Arnstein, D., Freedman, G., Dainer-Best, J., & Liss, A. (2011). Cognitive control in the intertrial interval: Evidence from EEG alpha power: Cognitive control in the intertrial interval. Psychophysiology, 48(5), 583–590. https://doi.org/10.1111/j.1469-8986.2010.01124.x

Delorme, A., & Makeig, S. (2004). EEGLAB: An open source toolbox for analysis of single-trial EEG dynamics including independent component analysis. Journal of Neuroscience Methods, 134(1), 9–21. https://doi.org/10.1016/j.jneumeth.2003.10.009

Enriquez-Geppert, S., Huster, R. J., Scharfenort, R., Mokom, Z. N., Zimmermann, J., & Herrmann, C. S. (2014). Modulation of frontal-midline theta by neurofeedback. Biological Psychology, 95, 59–69. https://doi.org/10.1016/j.biopsycho.2013.02.019

Foxe, J. J., & Snyder, A. C. (2011). The Role of Alpha-Band Brain Oscillations as a Sensory Suppression Mechanism during Selective Attention. Frontiers in Psychology, 2. https://doi.org/10.3389/fpsyg.2011.00154

Fries, P. (2015). Rhythms for Cognition: Communication through Coherence. Neuron, 88(1), 220–235. https://doi.org/10.1016/j.neuron.2015.09.034

Grabner, R. H., Stern, E., & Neubauer, A. C. (2003). When intelligence loses its impact: Neural efficiency during reasoning in a familiar area. International Journal of Psychophysiology, 49(2), 89–98. https://doi.org/10.1016/S0167-8760(03)00095-3

Gramfort, A., Luessi, M., Larson, E., Engemann, D. A., Strohmeier, D., Brodbeck, C., Parkkonen, L., & Hämäläinen, M. S. (2014). MNE software for processing MEG and EEG data. NeuroImage, 86, 446–460. https://doi.org/10.1016/j.neuroimage.2013.10.027

Hajihosseini, A., & Holroyd, C. B. (2013). Frontal midline theta and N200 amplitude reflect complementary information about expectancy and outcome evaluation: Frontal theta and N200 provide distinct information. Psychophysiology, 50(6), 550–562. https://doi.org/10.1111/psyp.12040

Händel, B. F., Haarmeier, T., & Jensen, O. (2011). Alpha Oscillations Correlate with the Successful Inhibition of Unattended Stimuli. Journal of Cognitive Neuroscience, 23(9), 2494–2502. https://doi.org/10.1162/jocn.2010.21557

Hanslmayr, S., Klimesch, W., Sauseng, P., Gruber, W., Doppelmayr, M., Freunberger, R., & Pecherstorfer, T. (2005). Visual discrimination performance is related to decreased alpha amplitude but increased phase locking. Neuroscience Letters, 375(1), 64–68. https://doi.org/10.1016/j.neulet.2004.10.092

Hanslmayr, S., Pastötter, B., Bäuml, K.-H., Gruber, S., Wimber, M., & Klimesch, W. (2008). The Electrophysiological Dynamics of Interference during the Stroop Task. Journal of Cognitive Neuroscience, 20(2), 215–225. https://doi.org/10.1162/jocn.2008.20020

Higgins, C., Liu, Y., Vidaurre, D., Kurth-Nelson, Z., Dolan, R., Behrens, T., & Woolrich, M. (2021). Replay bursts in humans coincide with activation of the default mode and parietal alpha networks. Neuron, 109(5), 882–893.e7. https://doi.org/10.1016/j.neuron.2020.12.007

Huycke, P., Verbeke, P., Boehler, C. N., & Verguts, T. (2021). Theta and alpha power across fast and slow timescales in cognitive control [Data set]. Zenodo. https://doi.org/10.5281/zenodo.4659714

Janssens, C., De Loof, E., Boehler, C. N., Pourtois, G., & Verguts, T. (2018). Occipital alpha power reveals fast attentional inhibition of incongruent distractors. Psychophysiology, 55(3), e13011. https://doi.org/10.1111/psyp.13011

Jasper, H. H. (1958). The ten-twenty electrode system of the International Federation. Electroencephalogr. Clin. Neurophysiol., 10, 370–375.

Jensen, O., & Mazaheri, A. (2010). Shaping Functional Architecture by Oscillatory Alpha Activity: Gating by Inhibition. Frontiers in Human Neuroscience, 4. https://doi.org/10.3389/fnhum.2010.00186

Jokisch, D., & Jensen, O. (2007). Modulation of Gamma and Alpha Activity during a Working Memory Task Engaging the Dorsal or Ventral Stream. Journal of Neuroscience, 27(12), 3244–3251. https://doi.org/10.1523/JNEUROSCI.5399-06.2007

Klimesch, W. (1999). EEG alpha and theta oscillations reflect cognitive and memory performance: A review and analysis. Brain Research Reviews, 29(2–3), 169–195. https://doi.org/10.1016/S0165-0173(98)00056-3

Kriegeskorte, N., Simmons, W. K., Bellgowan, P. S. F., & Baker, C. I. (2009). Circular analysis in systems neuroscience: The dangers of double dipping. Nature Neuroscience, 12(5), 535–540. https://doi.org/10.1038/nn.2303

Lee, T.-W., Girolami, M., & Sejnowski, T. J. (1999). Independent Component Analysis Using an Extended Infomax Algorithm for Mixed Subgaussian and Supergaussian Sources. Neural Computation, 11(2), 417–441. https://doi.org/10.1162/089976699300016719

Maris, E., & Oostenveld, R. (2007). Nonparametric statistical testing of EEG- and MEG-data. Journal of Neuroscience Methods, 164(1), 177–190. https://doi.org/10.1016/j.jneumeth.2007.03.024

Mazaheri, A., Nieuwenhuis, I. L. C., van Dijk, H., & Jensen, O. (2009). Prestimulus alpha and mu activity predicts failure to inhibit motor responses. Human Brain Mapping, 30(6), 1791–1800. https://doi.org/10.1002/hbm.20763

Narayanan, N. S., Cavanagh, J. F., Frank, M. J., & Laubach, M. (2013). Common medial frontal mechanisms of adaptive control in humans and rodents. Nature Neuroscience, 16(12), 1888–1895. https://doi.org/10.1038/nn.3549

Nigbur, R., Cohen, M. X., Ridderinkhof, K. R., & Stürmer, B. (2012). Theta Dynamics Reveal Domain-specific Control over Stimulus and Response Conflict. Journal of Cognitive Neuroscience, 24(5), 1264–1274. https://doi.org/10.1162/jocn_a_00128

O’Keefe, J. (1993). Hippocampus, theta, and spatial memory. Current Opinion in Neurobiology, 3(6), 917–924. https://doi.org/10.1016/0959-4388(93)90163-S

Peirce, J. W. (2007). PsychoPy—Psychophysics software in Python. Journal of Neuroscience Methods, 162(1–2), 8–13. https://doi.org/10.1016/j.jneumeth.2006.11.017

Peyrache, A., Khamassi, M., Benchenane, K., Wiener, S. I., & Battaglia, F. P. (2009). Replay of rule-learning related neural patterns in the prefrontal cortex during sleep. Nature Neuroscience, 12(7), 919–926. https://doi.org/10.1038/nn.2337

Pfurtscheller, G. (2003). Induced Oscillations in the Alpha Band: Functional Meaning. Epilepsia, 44(s12), 2–8. https://doi.org/10.1111/j.0013-9580.2003.12001.x

Phillips, J. M., Vinck, M., Everling, S., & Womelsdorf, T. (2014). A Long-Range Fronto-Parietal 5- to 10-Hz Network Predicts “Top-Down” Controlled Guidance in a Task-Switch Paradigm. Cerebral Cortex, 24(8), 1996–2008. https://doi.org/10.1093/cercor/bht050

Ridderinkhof, K. R. (2004). The Role of the Medial Frontal Cortex in Cognitive Control. Science, 306(5695), 443–447. https://doi.org/10.1126/science.1100301

Ritter, P., Moosmann, M., & Villringer, A. (2009). Rolandic alpha and beta EEG rhythms’ strengths are inversely related to fMRI-BOLD signal in primary somatosensory and motor cortex. Human Brain Mapping, 30(4), 1168–1187. https://doi.org/10.1002/hbm.20585

Ruge, H., & Wolfensteller, U. (2010). Rapid Formation of Pragmatic Rule Representations in the Human Brain during Instruction-Based Learning. Cerebral Cortex, 20(7), 1656–1667. https://doi.org/10.1093/cercor/bhp228

Sauseng, P., Klimesch, W., Doppelmayr, M., Pecherstorfer, T., Freunberger, R., & Hanslmayr, S. (2005). EEG alpha synchronization and functional coupling during top-down processing in a working memory task. Human Brain Mapping, 26(2), 148–155. https://doi.org/10.1002/hbm.20150

Sauseng, P., Tschentscher, N., & Biel, A. L. (2019). Be Prepared: Tune to FM-Theta for Cognitive Control. Trends in Neurosciences, 42(5), 307–309. https://doi.org/10.1016/j.tins.2019.02.006

Thut, G. (2006). Band Electroencephalographic Activity over Occipital Cortex Indexes Visuospatial Attention Bias and Predicts Visual Target Detection. Journal of Neuroscience, 26(37), 9494–9502. https://doi.org/10.1523/JNEUROSCI.0875-06.2006

van de Vijver, I., Ridderinkhof, K. R., & Cohen, M. X. (2011). Frontal Oscillatory Dynamics Predict Feedback Learning and Action Adjustment. Journal of Cognitive Neuroscience, 23(12), 4106–4121. https://doi.org/10.1162/jocn_a_00110

Verbeke, P., & Verguts, T. (2019). Learning to synchronize: How biological agents can couple neural task modules for dealing with the stability-plasticity dilemma. PLOS Computational Biology, 15(8), e1006604. https://doi.org/10.1371/journal.pcbi.1006604

Vertes, R. P., & Kocsis, B. (1997). Brainstem-diencephalo-septohippocampal systems controlling the theta rhythm of the hippocampus. Neuroscience, 4(81), 893–926.

Voloh, B., Valiante, T. A., Everling, S., & Womelsdorf, T. (2015). Theta–gamma coordination between anterior cingulate and prefrontal cortex indexes correct attention shifts. Proceedings of the National Academy of Sciences, 112(27), 8457–8462. https://doi.org/10.1073/pnas.1500438112

Wang, C., Trongnetrpunya, A., Samuel, I. B. H., Ding, M., & Kluger, B. M. (2016). Compensatory Neural Activity in Response to Cognitive Fatigue. The Journal of Neuroscience, 36(14), 3919–3924. https://doi.org/10.1523/JNEUROSCI.3652-15.2016

Winkler, I., Debener, S., Muller, K.-R., & Tangermann, M. (2015). On the influence of high-pass filtering on ICA-based artifact reduction in EEG-ERP. 2015 37th Annual International Conference of the IEEE Engineering in Medicine and Biology Society (EMBC), 4101–4105. https://doi.org/10.1109/EMBC.2015.7319296

Womelsdorf, T., Fries, P., Mitra, P. P., & Desimone, R. (2006). Gamma-band synchronization in visual cortex predicts speed of change detection. Nature, 439(7077), 733–736. https://doi.org/10.1038/nature04258

