## Supplementary Materials for "Theta and alpha power across fast and slow timescales in cognitive control"

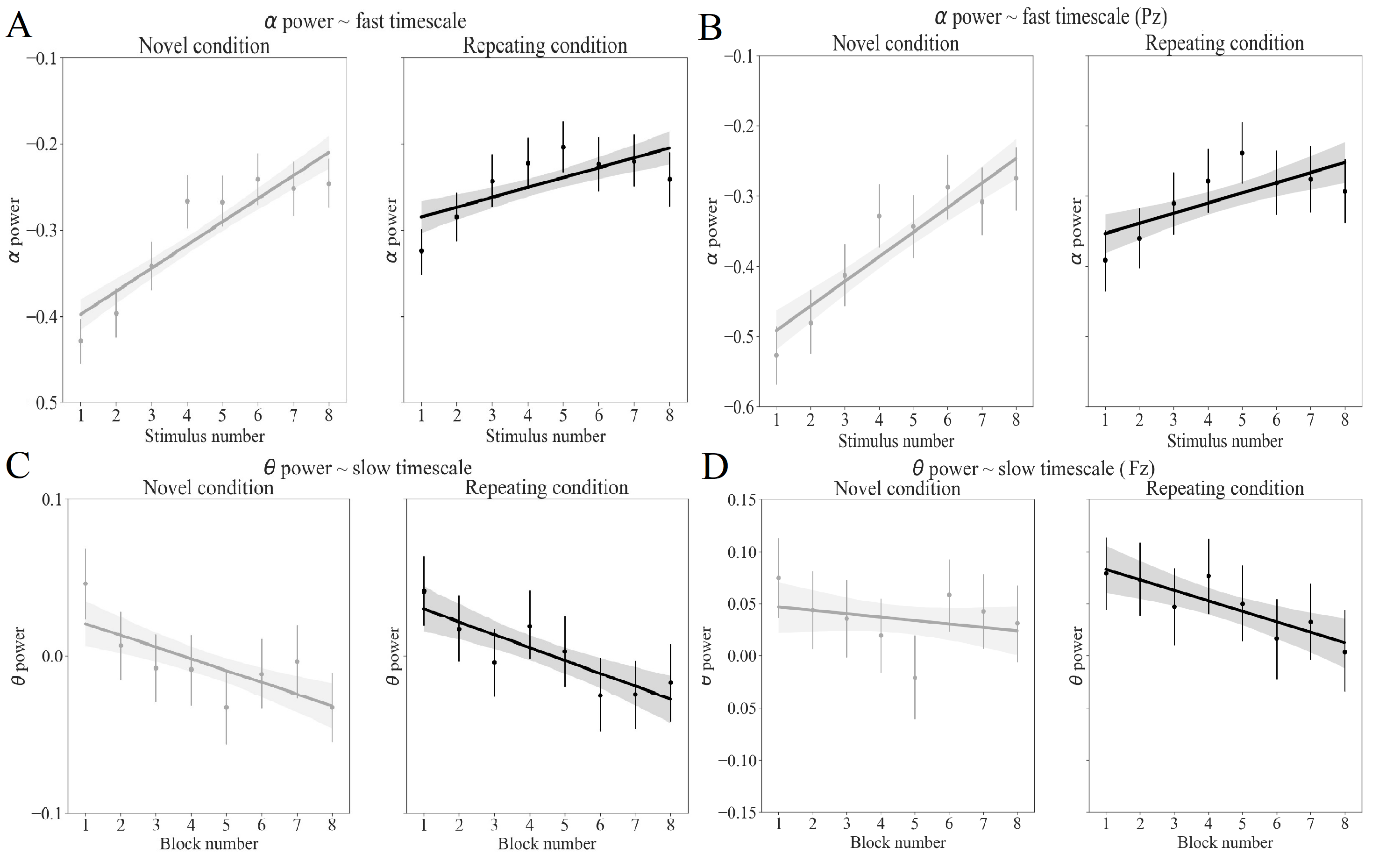
**Single-electrode alpha and theta**

**Figure S1**. Alpha and theta power when averaged across all electrodes and for one electrode. (A) This plot represents alpha power as a function of stimulus number across conditions. Subject-specific alpha is obtained by averaging the alpha power across all 64 electrodes. (B) This plot also represents alpha power as a function of stimulus number across conditions but now alpha is measured at the level of the single electrode Pz. Note that we chose to analyze alpha power at the level of Pz based on Figure 3B. We note that panels A and B look similar, suggesting that the observed alpha effects also occur at the level of an individual posterior electrode (C) Theta power as a function of block number and conditions. Theta power is obtained by averaging over all available electrodes. (D) Theta power as a function of block number and conditions, but now theta is obtained from one electrode Fz. We opted for Fz based on Figure 4B. We remark that panels C and D appear to show similar results which suggests that theta decreases both on average and on the level of a single frontal electrode.

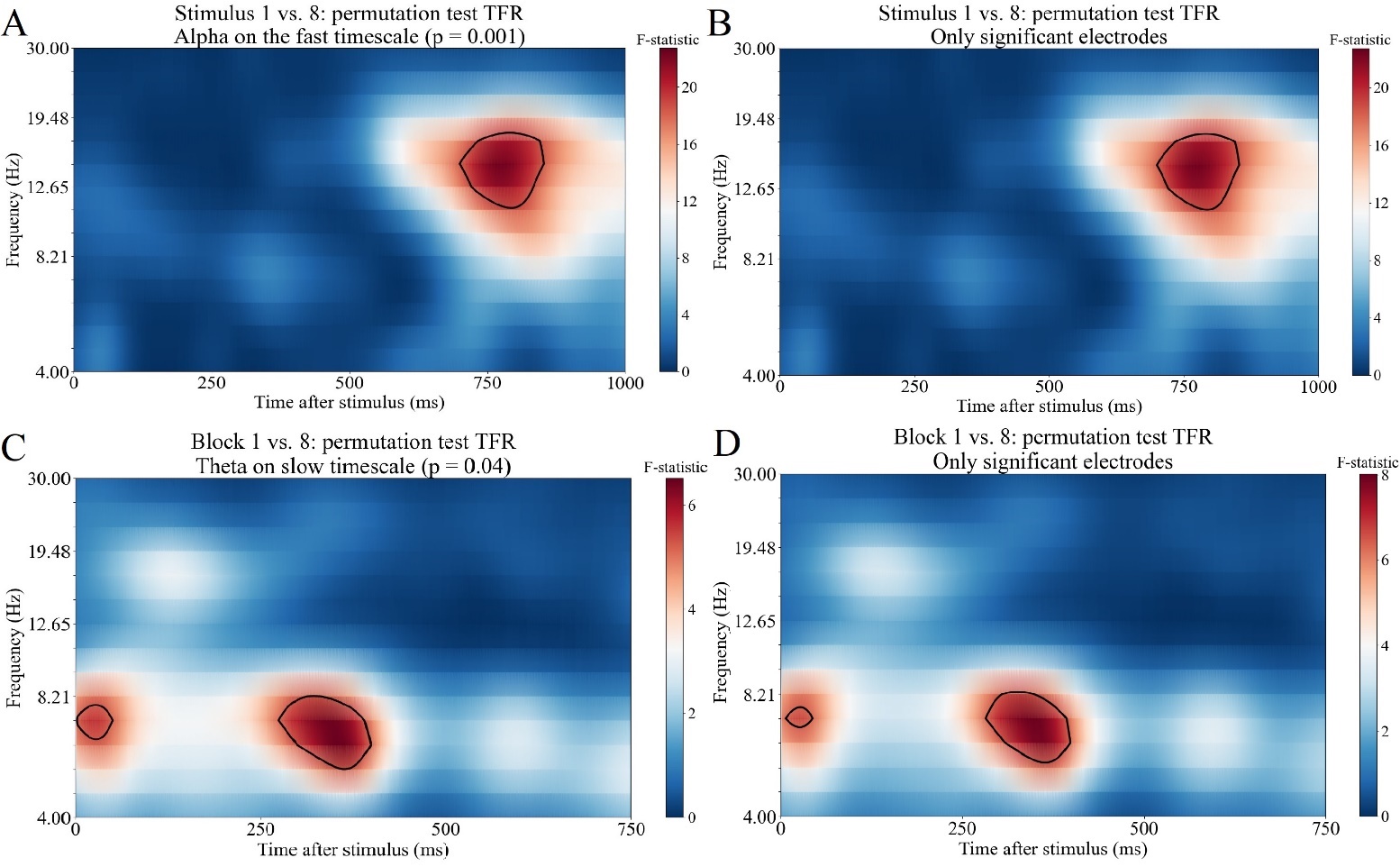
**All electrodes vs. a subset of significant electrodes**

**Figure S2.** In this figure we investigate which electrodes played a role in the significant clusters that we report. This was determined by revisiting the F values yielded after permutation testing (depicted in Figure 3A and 4A). First, we selected the time window of interest (700 – 850 ms and 250 – 400 ms for fast and slow timescale respectively) in this array of F values. Then we determined for each electrode separately whether this electrode has at least one data point in the frequency band of interest, namely alpha (8 – 12 Hz) for the fast timescale, and theta (4 – 8 Hz) for the slow timescale. When there is at least one electrode, we marked and retained it for further analyses. For the alpha cluster, we found that all 64 electrodes had a significant contribution. In contrast, for the theta cluster, we found that twelve out 64 electrodes (18.75%, all posterior electrodes) had no datapoints included in the cluster. In panel A we recreated the permutation time-frequency representation plot for the fast timescale (which is shown in Figure 3A). This panel can be compared with the permutation time-frequency representation plot created using only the significant electrodes, shown in panel B. Since all electrodes made a significant contribution on the fast timescale, panel B is identical to panel A. In panel C, we have recreated Figure 4A depicting the significant cluster found on the slow timescale. We can again compare this panel with the same plot created using only significant electrodes, which shows that plot is qualitatively the same, albeit it being based on only a subset of the total amount of electrodes. The main difference is that panel D contains more ‘extreme’ F-values as compared to panel C. This however is unsurprising since panel D is purely based on data from significant electrodes whereas panel C relies on data from all 64 electrodes.

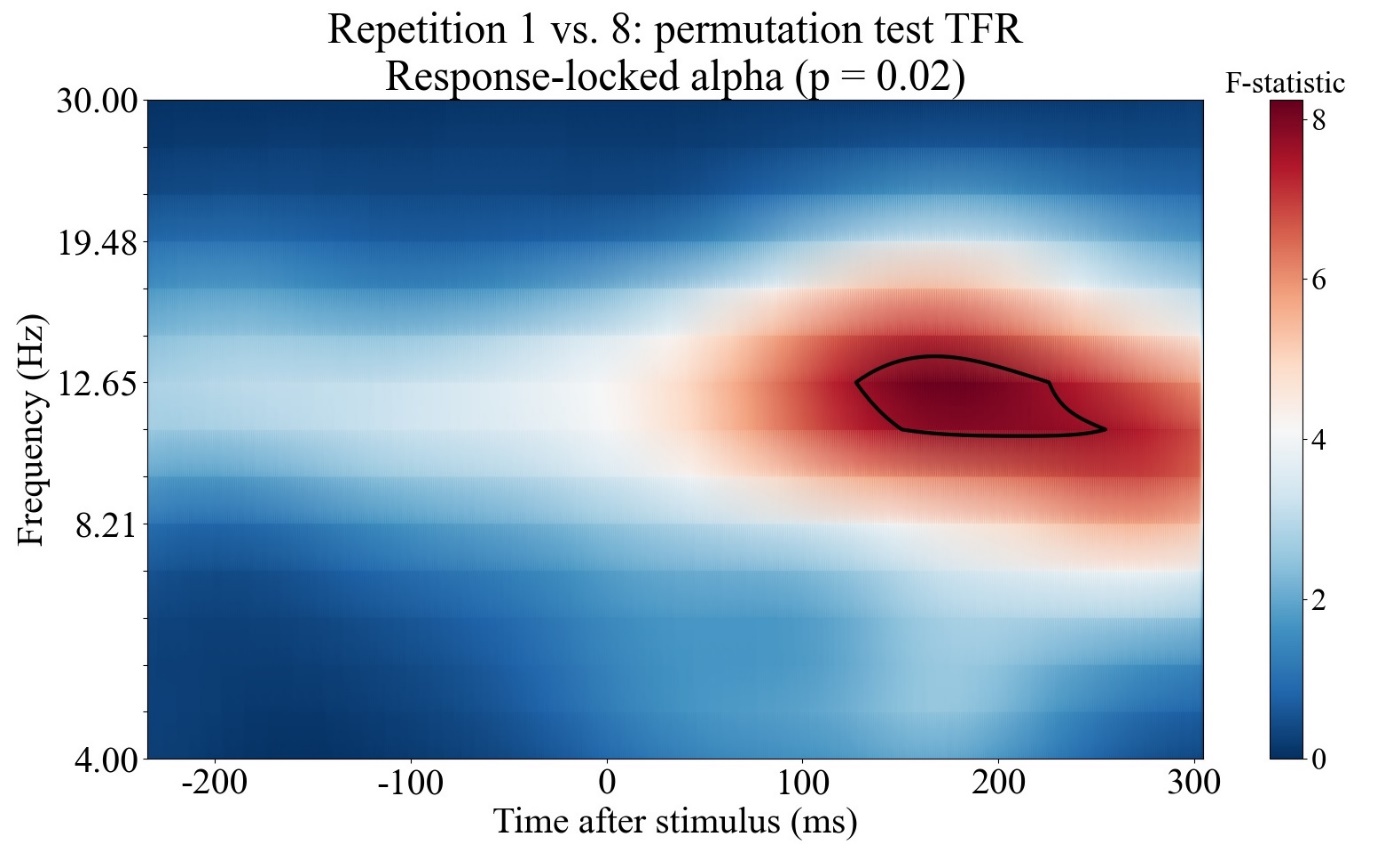
**Response-locked alpha band modulation**

**Figure S3**. This figure represents the significant cluster found in the response-locked data. This significant cluster occurs approximately 150 to 250 ms after stimulus onset, and appears around 10 to 14 Hz. The delineated area represents the 95% highest cluster statistics and only serves a visualization purpose. The cluster statistics are averaged across all 64 electrodes.
